# CRISPR/Cas9-mediated deletion of a kinetoplast-associated gene attenuates virulence in *Leishmania major* parasites

**DOI:** 10.1101/2024.03.11.584372

**Authors:** Fatemeh Darzi, Ali Khamesipour, Maryam Bahrami, Mahmoud Nateghi-Rostami

## Abstract

The CRISPR/Cas9 system has emerged as a powerful tool for precise genome editing, allowing for the deletion of genes, generation of point mutations, and addition of tags to endogenous genes. We employed an efficient CRISPR/Cas9 technique in *Leishmania major* to assess its efficiency in editing a kDNA-associated gene, universal minicircle sequence binding protein (UMSBP), which is involved in mitochondrial respiration and kinetoplast division.

We generated UMSBP C-tagged and *UMSBP* single knockout *L. major* (*Lm*UMSBP^+/−^) parasites using the CRISPR/Cas9 toolkit. C-tagged parasite were confirmed by PCR, flow cytometry and Western blot analyses. Gene expression of mitochondrial redox regulating enzymes, tryparedoxin peroxidase (TXNPx) and trypanothione synthetase (TryS), were analysed by real-time RT-PCR. Growth rate of promastigotes in culture and infectivity rate in macrophages were analysed *in vitro*. Mice were immunized by *Lm*UMSBP^+/−^ mutant strain and lesion size and parasite burden were measured upon challenge with live wild type (WT) *L. major*. Cytokines were titrated on supernatant of lymph nodes cell culture by sandwich ELISA.

Complete UMSBP deletion (*Lm*UMSBP^-^/^-^ null mutant) impaired promastigote survival, suggesting its essential role in parasite fitness. Despite this, we were able to produce attenuated *Lm*UMSBP^+/-^ parasites, which showed significant reduced growth in culture (*P*<0.05), increase in apoptosis (*P*<0.05) and downregulation of TXNPx and TryS gene expressions during growth of promastigotes compared to WT *L. major. Lm*UMSBP^+/-^ mutant strains did not cause ulcerative lesions in susceptible BALB/c mouse model. Furthermore, immunization of mice with *Lm*UMSBP^+/-^ parasites elicited a Th1 immune response with significantly high IFN-γ and low IL-4 production in cell culture (*P*<0.001) associated with partial protection against *L. major* WT challenge, as evidenced by reduced parasite burden and lesion development in BALB/c mice. Our findings demonstrate the potential of CRISPR/Cas9-edited *Lm*UMSBP^+/-^ parasites as live attenuated vaccine candidate against *Leishmania* infection.

**Author summary:** In this study, we utilized the powerful CRISPR/Cas9 technique to edit the genome of *Leishmania major*, a parasite responsible for causing leishmaniasis. Specifically, we targeted a gene called universal minicircle sequence binding protein (UMSBP), which plays a crucial role in the parasite’s mitochondrial function and replication. Using CRISPR/Cas9, we successfully created two types of parasites: one with a tagged UMSBP gene and another with the UMSBP gene completely knocked out. We produced an attenuated parasites with deleting UMSBP gene having reduced growth and increased apoptosis compared to wild-type parasites. Importantly, immunizing mice with these attenuated parasites induced a strong immune response, particularly IFN-γ secretion, and provided partial protection against infection with wild-type parasites. Our study suggests that CRISPR/Cas9-edited parasites could serve as promising live attenuated vaccine candidates against leishmaniasis.

## Introduction

Leishmaniasis is a neglected tropical disease which is caused by more than 30 species of *Leishmania* protozoan parasite. A wide range of clinical forms of the disease is described ranging from localized skin ulcers (cutaneous leishmaniasis, CL) to fatal systematic disease (visceral leishmaniasis, VL) (1). Worldwide an estimated of 1 billion people are at risk of infection in more than 100 countries. Currently there is no human vaccine available for any type of leishmaniases and various limitations of treatment including adverse effects and resistance emergence faced the management of the disease with severe difficulties (2).

Kinetoplast is a unique DNA structure (kDNA) in the single mitochondrion of trypanosomatids including *Leishmania* spp. kDNA composed of a network of circular DNA that are interlocked topologically viz: a few number of maxicircles (20–40 kb each) and a few thousand number of minicircles (0.5–10 kb each). Minicircles encode guide RNAs, function in the process of mRNA editing in most trypanosomatid species and contain two short sequences that are associated with replication initiation: a hexamer and a dodecamer, designated the universal minicircle sequence (UMS). These sequences have been mapped to the replication origins of the minicircle’s light (L) and heavy (H) strands, respectively. The UMS region is conserved in all members of trypanosomatid family. In addition, at the initiation of kDNA replication, binding activity of universal minicircle sequence binding protein (UMSBP), is alternatively regulated by mitochondrial redox regulating enzymes, tryparedoxin (TXN) and tryparedoxin peroxidase (TXNPx). Production of ATP is necessary for the proper functioning of the parasite, and the electron transport chain plays an important role in this process. A number of other mitochondrial genes produced by kDNA are involved in electron transport chain (3).

Despite decades of research devoted to develop a vaccine against leishmaniasis, such as first generation vaccines, recombinant antigens, DNA vaccination and live-attenuated strains, but we are still lacking a safe and effective vaccine for human leishmaniasis (4). Several of these candidate vaccines were shown to be safe and induce strong immune response in experimental models, but they have not provided satisfactory results in the efficacy trials. Therefore, the search for a new prophylactic vaccine candidate for management of leishmaniasis should be a global priority. Live-attenuated *Leishmania* vaccine candidates are promising alternatives since they have the advantage of the better efficacy to induce long-term protective immune responses as compared to killed parasite vaccines, but without the risk of pathology and disease development as in live vaccines (5, 6). In live attenuated vaccines, parasites are used which turned nonpathogenic by different procedures such as biological and physicochemical methods or gene knockout approach. So far, several genes of the *Leishmania* parasite such as *centrin, gp63, CPs, P27* and other genes have been used as possible target for production of attenuated strains (7, 8). The discovery of CRISPR/Cas based gene editing allowed genetic manipulation of parasites and/or vectors for disease control and selection of safer mutant live-attenuated *Leishmania* parasites to investigate as vaccine candidate (9, 10).

In this study, we have generated a UMSBP C-tagged strain as well as a UMSBP-knock out attenuated strain of *L. major* by an efficient and feasible CRISPR/Cas9 method. These strains could be used to further study the organization and biological function of the kinetoplast and its related structures. Furthermore, since kDNA is a unique structure in trypanosomatids and due to the role of kDNA in the electron transfer chain and ATP generation of the cell, it is possible that deletion of this gene produces a non-pathogenic strain suitable as a candidate vaccine against leishmaniasis.

## Materials and methods

### 1. *Leishmania* strain and culture

*L. major* Friedlin (MHOM/SU/73/5-ASKH) strain (1×10^5^ logarithmic promastigotes per mL) were cultured at 26 °C in M199 medium (Gibco, USA) (pH 7.4) supplemented with 15% heat-inactivated fetal calf serum (FCS) (in case of medium which was used after transfectin, 20% FCS), 100 U/mL penicillin/100 μg/mL streptomycin, 40 mM HEPES (4-(2-hydroxyethyl)-1-piperazineethanesulfonic acid), 5 µg/mL hemin, 10 µg/mL folic acid, 8 µM 6-biopterin,1 mg/L biotin and 1x RPMI 1640 vitamins solutions (Sigma, USA). Parasites in culture were passaged to fresh medium at a one 20th-50th dilution once a week.

Relevant selective drugs were added to the medium at the following concentrations: 200 μg/mL hygromycin B (Invitrogen, USA), 30 μg/mL puromycin dihydrochloride (Sigma), and 40 μg/mL blasticidin S hydrochloride (Invitrogen).

### 2. CRISPR/Cas9 mediated deletion and tagging of UMSBP gene

In this CRISPR/Cas9 strategy, pPLOT and pT plasmids are used which serve as template DNA for PCR amplification of repair cassettes for genome editing by homologous recombination. The amplicons are transfected together with the relevant sgRNA templates into a kinetoplastid cell line expressing Cas9 nuclease and T7 RNA polymerase (T7 RNAP) for *in vivo* transcription of sgRNAs. Specific plasmids developed for the CRISPR system in *Leishmania* were a kind gift from Institut Pasteur de Paris (Unité de Parasitologie Moléculaire et Signalisation Département de Parasitologie et Mycologie).

The information related to the gene and its protein product was extracted from various databases such as NCBI, Ebi, SWISSPROT and TriTrypDB. Different sequences were aligned with each other with Clustal to identify conserved loci and divergent parts.

### 2.1. Generation of *L. major* cell line expressing Cas9 and T7 RNAP

The pTB007 plasmid used contained the genes encoding the *Streptococcus pyogenes* Cas9 nuclease and the T7 RNA-polymerase (Cas9/T7), along with the hygromycin resistance gene. Ten µg of pTB007 was transfected into 4×10^7^ mid logarithmic phase *L*. *major* promastigotes in 83 μl 3× Tb-BSF transfection buffer (200 mM Na2HPO4, 70 mM NaH2PO4, 15 mM KCl, 150 mM HEPES, pH 7.4) (11) mixed with 25 μl 1.5 mM CaCl2 and 42 μl ddH2O using 2 mm gap cuvettes with 3.5 msec/500 volt in BIORAD Gene Pulser Xcell Electroporation System (BioRad, USA). Transfected cells were immediately transferred into 5 mL prewarmed medium in 25 cm^2^ flasks and left to recover overnight at 26 °C. Transfected cells were then selected for 200 μg/mL hygromycin B (Invitrogen) resistance following 3 consecutive passages. The live promastigote cell line which expressed the Cas9 enzyme and T7 RNA-polymerase were used for next steps.

#### 2.2. Design of the sgRNAs and the donor DNA primers

The sgRNA primers were composed of a T7 promoter sequence in 5’ end (GAAATTAATACGACTCACTATAGG), a homology sequence with G00 Primer (Cas9-backbone-start) in 3’ end (GTTTTAGAGCTAGAAATAGC) and a sgRNA target sequence.

The sgRNA target sequences used to delete and c-tagging the *Lm*UMSBP gene were obtained from LeishGEdit.net. The sgRNAs had the highest-scoring 20 nt sequence within 105 bp of UTRs bordering upstream and downstream of the target locus. Sequences followed by a Protospacer Adjacent Motif (PAM) sequence in 3’ end (http://www.leishgedit.net). The sequences of the sgRNAs were blasted against the *L*. *major* genome in TriTrypDB to verify that the sgRNAs were specific for *Lm*UMSBP exclusively (*E* value = 0.001 and 8e^-5^).

The upstream and downstream donor DNA primers were composed of a homology flank sequence (HFN30) of 30 nucleotides which was selected among the 300 nucleotides closest to the target locus in UTRs as well as a binding site homologue to the plasmid templates (PT or pPLOT) (Table 1). The repair cassette for tagging had mNeonGreen fragment as fluorescence tag.

**Table 1.**
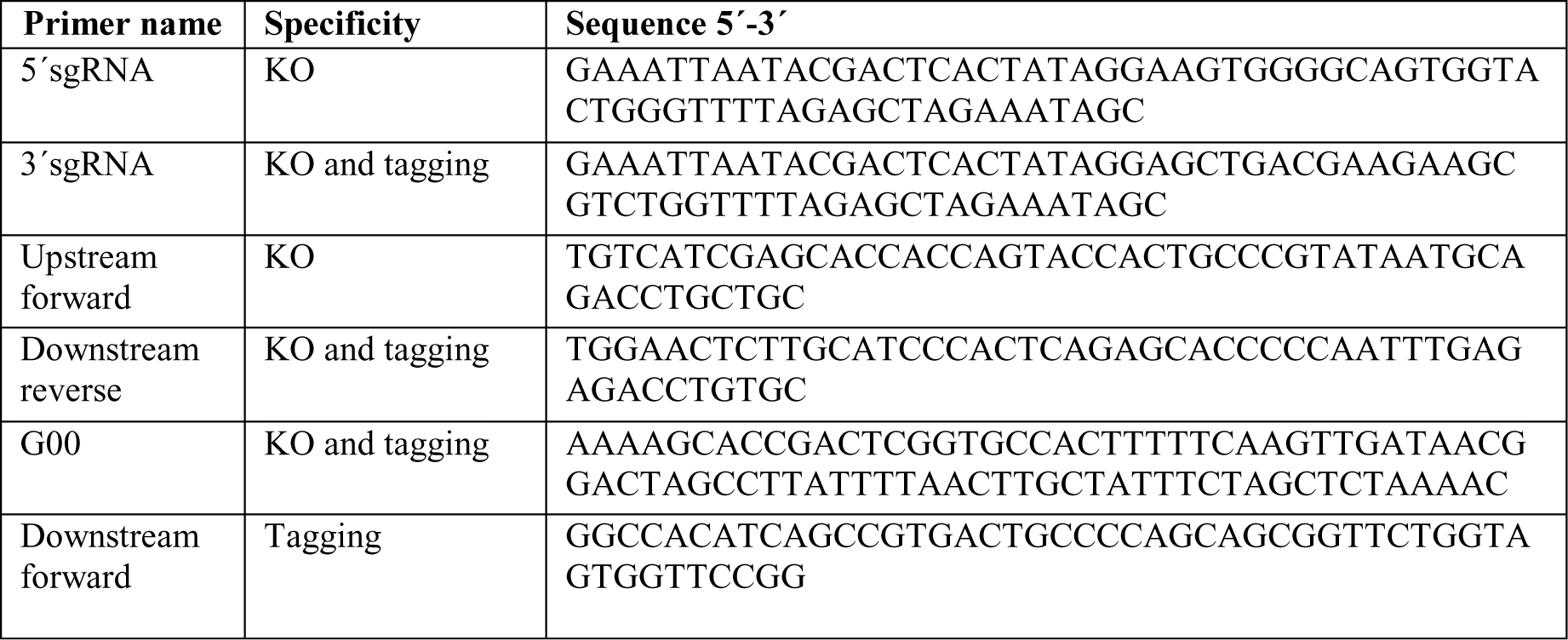
Sequence of primers used in CRISPR/Cas9 experiment.

The repair cassette showed an E value of 5e^-9^ suggesting very high specificity of the system. The sgRNA target sequences and the homology arms on the repair cassette fully matched the target sequence of *Lm*UMSBP.

#### 2.3. Generation of sgRNA templates and donor DNA constructs

DNA fragments encoding *Lm*UMSBP specific 5’ and 3’ guide RNAs for upstream and downstream to the target gene were PCR amplified. All primers are listed in Table 1.

In order to amplify the sgRNA templates, the reaction mixture consisted of 2 µM of 5’ or 3’ guide RNAs, 2 µM primer G00 fragment, 200 µM dNTPs mix (Bio Basic, Toronto, Canada), 1X EHF buffer with 15 mM MgCl2 (Roche), 1 U HiFi enzyme mix (Expand High Fidelity PCR System, Roche, USA) and 1.5 mM MgCl2 (Roche, Basel, Switzerland). The PCR conditions included an initial denaturation at 98 °C for 30 sec, followed by 35 cycles of 98 °C for 10 s, 60 °C for 30 s and 72 °C for 15 s, and a final extension step of 72 °C for 7 min. To assess the presence of the expected product, 2 *μ*L of the 20 μL reaction was run on a 1% agarose gel. All PCR products were heat-sterilized at 95 °C for 5 min before transfection.

For the PCR amplification of the targeting fragments of pPLOT and pT cassettes, the reaction mixture consisted of 2 μM of donor primers mix, 200 µM dNTPs (Bio Basic), 15 ng circular pPLOT or pT plasmid, 3% (v/v) DMSO (Dimethyl sulfoxide; Sigma Aldrich), 1 U HiFi enzyme mix (Roche), 1X buffer (Roche) and 1.875 mM MgCl2 in a total volume of 40 μL. The PCR conditions included initial denaturation at 94 °C for 5 min followed by 40 cycles of 94 °C for 30 s, 65 °C for 30 s and 72 °C for 2 min 15 s, and lastly the final extension at 72 °C for 7 min. The presence of the expected product was assessed by running 2 μL of the 40 μL reaction on a 1% agarose gel. All PCR products were heat-sterilized at 95 °C for 5 min before transfection.

For tagging strategy, the donor DNA construct was generated by the PCR amplification of a cassette with puromycin resistant marker from a pPLOT plasmid (pPLOT puro-mNeonGreen-puro) with downstream donor DNA primers for C-terminal tagging. (mNeonGreen :: Myc).

For knockout strategy, two donor DNA constructs were generated by PCR amplifications of two resistance cassettes from pT plasmids (pTBlast and pTPuro) with upstream forward and downstream reverse donor DNA primers.

The resulting fragments promote the integration of the drug resistance cassette by homologous recombination for repair of the DSB (double-strand break) at the target site.

#### 2.4. Production of *Lm*UMSBP KO and c-terminal tagging mutants

To generate *Lm*UMSBP c-tagging mutants we designed a 3’ sgRNAs to create DSB downstream of the *Lm*UMSBP coding region with a repair cassette containing puromycin resistance marker. To generate *Lm*UMSBP KO mutants we designed two 5’ and 3’ sgRNAs to create DSB upstream and downstream of the *Lm*UMSBP coding region with two repair cassettes containing puromycin and blasticidin S resistance markers.

The PCR amplified products (4 μg of each), including pooled sgRNA template(s) together with the donor DNA construct(s), were transfected into mid-log phase cell line expressing Cas9 enzyme and T7 RNA-polymerase as described in previous section. Transfected cells were further selected for resistance to 200 μg/mL hygromycin B (Invitrogen), 20 μg/mL puromycin dihydrochloride (Sigma), and 40 μg/mL blasticidin S hydrochloride (Invitrogen).

### 3. Diagnostic PCRs to confirm tagging and deletion of *Lm*UMSBP gene

To assess the loss of the target gene in the KO and tagging strains, genomic DNA was isolated from parasites collected after two passages post-transfection with the DNeasy Blood & Tissue Kit (Qiagen Hilden, Germany) and analyzed using specific primers (Table 2). The forward primer located on the coding region of the gene and the reverse primer was located on the antibiotic resistance gene (Figure1).

**Table 2.**
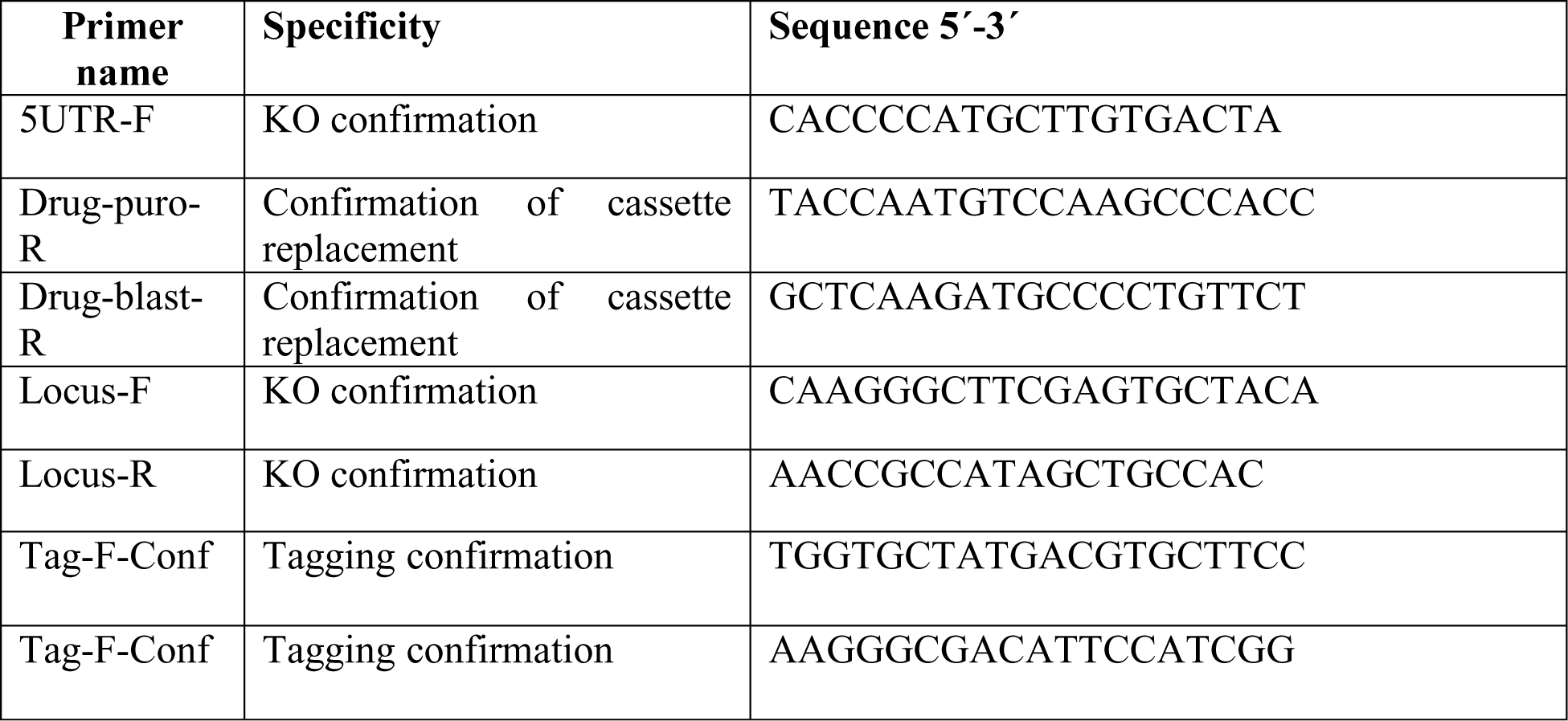
Sequence of primers used in KO and tagging confirmation procedures.

**Fig. 1.**
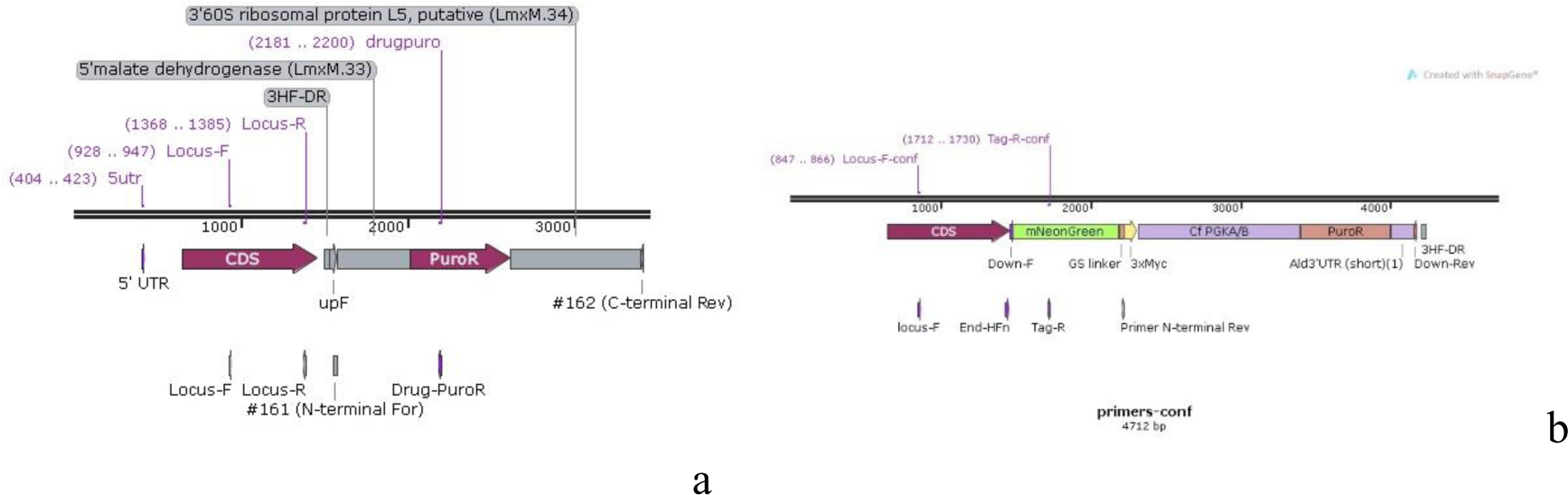
Schematic map of UMSBP gene locus, insertion cassette and primers binding sites of KO (a) and tagged (b) mutant *L. major* strains.

The reaction mixture consisted of 0.5 μM of each of primers, gDNA (30 ng), dNTPs (0.2 mM) (Biobasic, Markham, Canada), 1 U HiFi enzyme mix (Roche), 1X buffer, 1.5 mM MgCl2 and 2 mM MgSO4 (Bio Basic) in a total volume of 20 μL. PCR conditions included initial denaturation at 94 °C for 5 min followed by 35 cycles of 94 °C for 30 s, 60 °C for 30 s, and 72 °C for 2 min 15 s and a final extension at 72 °C for 7 min. Five microliters of PCR products was run on a 1% agarose gels.

### 4. Growth analysis of promastigotes

*L*. *major* wild type (WT*Lm*), Cas9 /T7 RNAP expression pTB007 cell line (pTB007 CLine), and the *Lm*UMSBP mutant cells (KO) were cultured as promastigotes at 26 °C in M199 medium with supplements (above). Cells were seeded at a concentration of 5×10^5^ cells/mL, and counted daily during 10 consecutive days. The curves were obtained from three independent repeats.

### 5. Sodium dodecyl-sulfate polyacrylamide gel electrophoresis (SDS-PAGE) and western blot analysis

About 5×10^7^ promastigotes in the mid-log phase of growth were harvested from culture (10 mL) and washed twice with PBS, pH 7.4. The cell pellet was resuspended in RIPA lysis buffer containing 150 mM NaCl, 25 mM Tris HCl, pH 7.6, 1% NP-40, 1% sodium deoxycholate, 0.1% SDS, pH 7.6, and protease inhibitor cocktail (Roche) and supplemented with 5 mM sodium orthovanadate and 1% Triton X-100.

Cells were resuspended in 100 μl of 2X Laemmli SDS sample buffer (63 mM Tris-HCl, 10% glycerol, 2% SDS, 0.0025% bromophenol blue, pH 6.8) and heated at 95 °C for 5 min. A total of 20 µg protein (40 μL) was denatured and separated by 10% SDS-PAGE for 1–2 h at 100 V., and blotted onto polyvinylidene difluoride (PVDF) membranes, and probed using specific primary and secondary antibodies. To this end, anti-Flag M2 monoclonal antibody with 1/1000 dilution (Sigma, F3165) was used. Anti-mouse IgG–Peroxidase antibody was used with 1/500 dilution (Sigma, A9044). For detecting tag fragment, primary antibody was anti-Myc with 1:1000 dilution (Biosensis, R-1319-100). As the secondary antibody, peroxidase-labelled goat anti-rabbit IgG antibody was used with 1/500 dilution (Thermo Scientific, 31462). 3,3’ Diaminobenzidine (DAB) substrate solution in hydrogen peroxide was used on the membrane until the desired development is achieved.

### 6. Flow cytometry analysis

Tagged *L. major* promastigotes were seeded at 1×10^5^ promastigotes/mL and cultured for 10 days, and aliquots were taken every 24 h for analysis. Cell viability was verified by incubation of the cells with 20 μg/mL propidium iodide (PI) for 30 min. The stained cells were analyzed using the ImageStream X Mark II Imaging flow cytometer (Merck Millipore, USA) with an X60/0.9 objective. Data from channels representing bright field, and fluorescence (PI) emission at 488 nm (to evaluate cell viability) were recorded for 20,000 cells for each analyzed sample. IDEAS software generated the quantitative measurements of the focused, single live cells for all examined cell strains. Cell shape was quantified using circularity and length features applied to the bright field image processed by an Adaptive Erode mask. Representative scatter plots are shown for single cells and for circularity (cell shape). Recorded emission of the PI in the gated population evaluated cell viability.

### 7. Fluorescence Microscopy

*L. major* promastigotes expressing the fluorescent fusion protein UMSBP-mNG-myc were imaged by fluorescence microscopy. Briefly, parasites were harvested from logarithmic phase (day 2-3) culture at the rate of 2×10^6^/mL, washed three times in PBS by centrifugation at 800 x *g* for 5 min and fixed with absolute ethanol and stained with DAPI at 1 *μ*g/mL final concentration for 15 min. The cells were analysed with a NPL Fluotar 50x/1.00 Oil, 160/- oil immersion objective lens on a Leitz LaborLux 12 Fluorescence Microscope (Leitz, Germany).

### 8. Assessment of apoptosis markers

Apoptosis was measured by the annexin V kit (Thermo Fisher Scientific, USA), which measures the binding of phosphatidylserine to annexin V in the early stages of apoptosis, late stages and necrosis, in WT*Lm*, pTB007 CLine and KO strains. The cells were harvested at 1×10^6^, washed with PBS, pH 7.2, and resuspended in 1x binding buffer. Then, 5 μL of FITC-conjugated Annexin V and 1 µL of the 100 µg/mL PI working solution were added to 100 μL of the cell suspension. Cells were incubated 15 min at room temperature in dark. Then 0.5 mL of 1X binding buffer was added, mixed and kept on ice then analyzed by flow cytometry (BD FACS Calibur, USA) measuring the fluorescence emission at 520 nm (FL1, Anexin V-FITC) and > 617 nm (FL3-PI). Cell viability was verified by incubation of the cells with 20 μg/mL PI for 30 min. The stained cells were analyzed using the ImageStream X Mark II Imaging flow cytometer (Merck Millipore) with an X60/0.9 objective. Data from channels representing bright field, and fluorescence (PI) emission at 488 nm (to evaluate cell viability) were recorded for a minimun of 20,000 cells for each analyzed sample.

## 9. *In vitro* infection assay of mice macrophages

*L. major* UMSBP KO, pTB007 CLine, and WT*Lm* strains, were seeded and allowed to grow for 5 days to reach stationary growth phase. Cells were washed with RPMI 1640 counted and used to infect J774 macrophages. Briefly, 2×10^5^ macrophages were pre-seeded a day in advance in 8-wells chamber slide (Nunc, Lab-Tek, USA,) and allowed to adhere, unattached cells were washed. *Leishmania* promastigotes were added at a ratio of 10:1 (parasite:macrophage) in 200 μl RPMI 1640 medium, the cells were washed with PBS, pH 7.4, to remove extracellular parasites, and incubated at 37 °C with 5% CO2 for 72 h The infected macrophages were fixed in absolute methanol, stained with Giemsa, and examined microscopically and the number of infected cells and internalized parasites were counted in 100 macrophages, at 6, 24 and 72 h after infection. The infectivity index was calculated by multiplying the number of infected macrophages by the number of amastigotes per infected macrophage.

### 10. Real time RT-PCR

The promastigotes from stationary phase (day 5-6 of culture) were harvested and used for RNA extraction followed by reverse transcription of mRNA to cDNA. The cDNA was then used as template for real-time RT-PCR using specific primer targeting tryparedoxin peroxidase (TXNPx) and trypanothione synthetase (TryS), as previously described (12).

The cell pellet was resuspended in cold PBS, pH 7.2, and added with 1 mL of TRIzol (Sigma) per 1×10^6^ cells. RNA was solubilized through pipetting, then, 0.2 mL chloroform was added per 1 mL of homogenate, followed by centrifugation for 15 min, 12,000 × *g* at 4 °C. The upper phase was collected, 0.5 mL of isopropanol per 1 mL of TRIzol was added, centrifuged at 12,000 x *g*, 4 °C for 10 min, the supernatant was discarded, and the RNA pellet was washed with 1 mL 75% ethanol at 7,500 x *g* for 5 min. The pellet was dissolved in DEPC-treated water and incubated on heat block at 55-60 °C for 10 min. The purity of RNA samples was assessed by the ratio of optical densities (ODs) at 260/280 nm using UV spectrophotometry (Thermo Fisher Scientific, Waltham, MA, USA).

Reverse transcription was carried out using AllScript Reverse Transcriptase kit with RevertAid MMuLV enzyme (Biotechrabbit, Hennigsdorf, Germany). Briefly, in a 20 μL reaction mixture, 2 μL of oligo (dT)12-18 primer (1 μM), 1 μL of dNTP mix (0.5 mM of each), 2 μg of total RNA, 0.5 μL of RNase inhibitor (1 U/μL) and 4 μL of 5x buffer mixed and incubated at 50 °C for 30 min. The enzyme was inactivated by heating at 95 °C for 5 min, and product was proceeded to the next step.

All reactions were performed in a total volume of 20 μL which contained 2 μL cDNA sample, 10 μL 2x SYBR Green PCR Master Mix (Qiagen) and 0.5 μL (0.5 μM each) of each primer targeting TryS (F: 5’-ggtactacgacgaagcca-3’ and R: 5’-tgatccgcgacacatacgta-3’) and TXNPx (F: 5’-ccatactactgcgacggcga-3’ and R: 5’-gcccacagcaccttcattcg-3) and 7.5 μL dH2O. For normalizing, the housekeeping gene glyceraldehyde-3-phosphate dehydrogenase (GAPDH) was used as the internal standard. Three technical replicates were performed for each experiment. A no-template control was included for each gene. Two-step thermal profile as a PCR program was set up on the Rotor-Gene 6000 Series Software (version 1.7) of Rotor-Gene machine (Corbett Life Science, USA) with the following thermal cycling profile: 95 °C for 5 minutes, followed by 45 cycles at 95 °C for 60 seconds and 60 °C for 45 s. The amplification process was followed by melting curve analysis. The threshold cycle (Ct) of each sample was used in calculation of relative gene expression using the 2^−ΔΔCt^ method as described (13).

### 11. Mice infection, immunization, and live challenge

For safety studies, two groups (10 animals per group) of female 6 to 8 week-old BALB/c mice were inoculated subcutaneously (s.c.) in the footpad with 1 × 10^7^ *L. major* WT or KO promastigotes in the stationary phase. Footpad thickness was measured weekly after infection using a metric caliper.

For vaccination and immunological studies, two groups of BALB/c mice (10 mice/group) were immunized intradermally (i.d.) in the left ear with either 2×10^6^ KO parasite in the stationary phase, or PBS in a volume of 20 μl. After 4 weeks (day 28) post immunization, both groups of mice were challenged i.d. in the contralateral ear with 2×10^6^ live stationary phase (day 5-6 of culture) *L. major* promastigotes in 50 μl volume. The progress of infection was recorded by weekly measurement of ear lesion size using a metric caliper.

### 12. Determination of parasite burden

For *in vivo* safety studies, 8 weeks post infection and for efficacy studies, 10 weeks post challenge three mice of each group were sacrificed and the spleen and draining lymph nodes were extracted to perform parasite burden by limiting dilution assay (LDA) as previously described (14). Briefly, tissues were aseptically removed, weighed and washed with sterile PBS, pH 7.2, then homogenized using a homogenizing pestle in Schneider’s *Drosophila* Medium (Gibco) supplemented with 20% heat-inactivated FCS (Gibco) and 1% Penicillin/Streptomycin. The homogenate was diluted in eight serial 10-fold dilutions with the same medium (from 1:1 to 1:10,000,000 in a total volume of 1.8 ml). About 100 μl of each suspension were distributed to 12 replicate wells of 96-well microtiter plates, which were covered to prevent medium evaporation and external contamination. The plates were incubated at 25 ±1 °C for 10 days. Positive (the presence of motile parasite) and negative (the absence of motile parasite) wells were specified using an inverted microscope (Zeiss, Germany). The parasite burden was determined after logarithmic calculation of the microscopic results by ELIDA software (15) and presented as the number of parasites/mg of infected tissues.

### 13. Soluble *Leishmania* antigen (SLA) preparation

Stationary phase promastigotes of *Leishmania major* (MRHO/IR/75/ER) strain were harvested at day 5 of culture from M199 medium supplemented with 10% FCS (Gibco), washed 3 times in cold sterile PBS, pH 7.2, and used for preparation of SLA as previously described by (16). Briefly, protease inhibitor cocktail enzyme (Sigma) was added at 100 µL per 1×10^9^ promastigotes, then the parasites were disrupted by 10 cycles of freeze (liquid nitrogen at −196 °C)-thawing (water bath at 37 °C) followed by sonication with two 20-second blasts at 4 °C. Parasite suspension was centrifuged at 30,000 x *g* for 20 min, and the supernatant was collected and re-centrifuged at 100,000 x *g* for 4 h. SLA protein concentration was measured by Bradford protein assay. Finally, the supernatant was sterilized using 0.45 µm membrane filter and stored at −20 °C until use.

### 14. Cell culture and ELISA cytokine assays

Four weeks after immunization, mice were sacrificed and the draining lymph nodes were extracted and homogenized with a homogenizing pestle. Lymph node suspensions were then plated at a density of 2×10^6^/mL in 96-well plates in RPMI 1640 medium supplemented with 10% FCS, 10 mM/L HEPES, 2 mM L-glutamine, 100 U/mL penicillin G and 100 mg/mL streptomycin (Gibco Invitrogen) in the presence of either 10 µg/mL lipopolysaccharide (LPS), 25 µg/mL of SLA or without stimulation. Plates were incubated at 37 °C with 5% CO2 in humidified atmosphere, and the levels of IL-12, IFN-γ, IL-4 and IL-10 were titrated on for 72-h culture supernatants using sandwich ELISA method (R & D System, Minneapolis, MN, USA) as previously described (17). Briefly, the plates were coated with anti-IL-12/IFN-γ/IL-4/IL-10 mAb overnight, and after blocking the wells using PBS plus 0.1% (w/v) bovine serum albumin (BSA), 50 µL of culture supernatants or standard dilutions were added. Then, biotin-labelled mAb, streptavidin– horseradish peroxidase (HRP) and 3,3’,5,5’-tetramethyl benzidine (TMB) substrate were added and the reaction was stopped with 2 N H2SO4 solution. The plates were washed after each step of incubation using PBS+0.05% (v/v) Tween-20. The plates were read at 450 nm using a reader (BioTek, Winooski, VT, USA), and the mean ODs of triplicate samples were compared with the standard curves prepared using recombinant cytokines.

### 15. Statistical analysis

Student’s *t*-test and repeated measure ANOVA tests were used to compare between groups. For some comparisons, nonparametric tests were chosen because the samples did not follow a Gaussian distribution. Nonparametric tests of Mann–Whitney, Kruskal-Wallis and Dunn’s post-test for paired comparisons were used for statistical analysis of the data using SPSS statistics version 20 (SPSS Inc, Chicago, IL, USA) and GraphPad Prism version 8.01 (GraphPad Software Inc, La Jolla, CA, USA) softwares. *P* value of <0.05 was considered to be significant. *P* value of ≤0.05 was considered to be statistically significant.

## Results

### 1. Bioinformatics data

It is crucial to know the exact sgRNA sequence and protospacer-adjacent motif (PAM) to successfully tag or delete genes in *Leishmania* spp. using CRISPR/Cas9. Considering the possibility of nucleotide diversity in the regions related to gRNA binding, it was also necessary to compare the different sequences related to the gene and its upstream and downstream regions by blast and identify the conserved regions. This review was necessary before gRNA design. In addition, by comparing this gene between different strains, it was possible to perform a similar method to remove the desired gene in other strains. We determined that UMSBP has conserved C-terminal extension whereas it has non-conserved N-terminal in comparison of other *Leishmania* spp. We used *L. major* reference genome to design the primers required to generate the donor DNA and the corresponding sgRNAs. Sequences were identified using the EuPaGDT CRISPR gRNA Design Tool with similar target parameters as those used in the LeishGEdit strategy.

### 2. Expression of Cas9 and T7 RNA polymerase did not affect parasite growth and differentiation

We first transfected *L. major* promastigotes with the plasmid pTB007 expressing (i) the *Streptococcus pyogenes* Cas9 nuclease gene (hSpCas9) with a nuclear localization signal and three copies of the FLAG epitope at the N-terminus, (ii) T7 RNA polymerase (T7 RNAP) and (iii), a hygromycin resistance gene (18). We confirmed the expression of Cas9 by Western blot analysis and showed that this expression did not alter the growth of *L. major* promastigotes (CLine) (Figure 2). We found that Cas9 is also expressed in axenic amastigotes The presence of pTB007 did not interfere with axenic amastigote differentiation or proliferation (Data not shown).

**Fig. 2.**
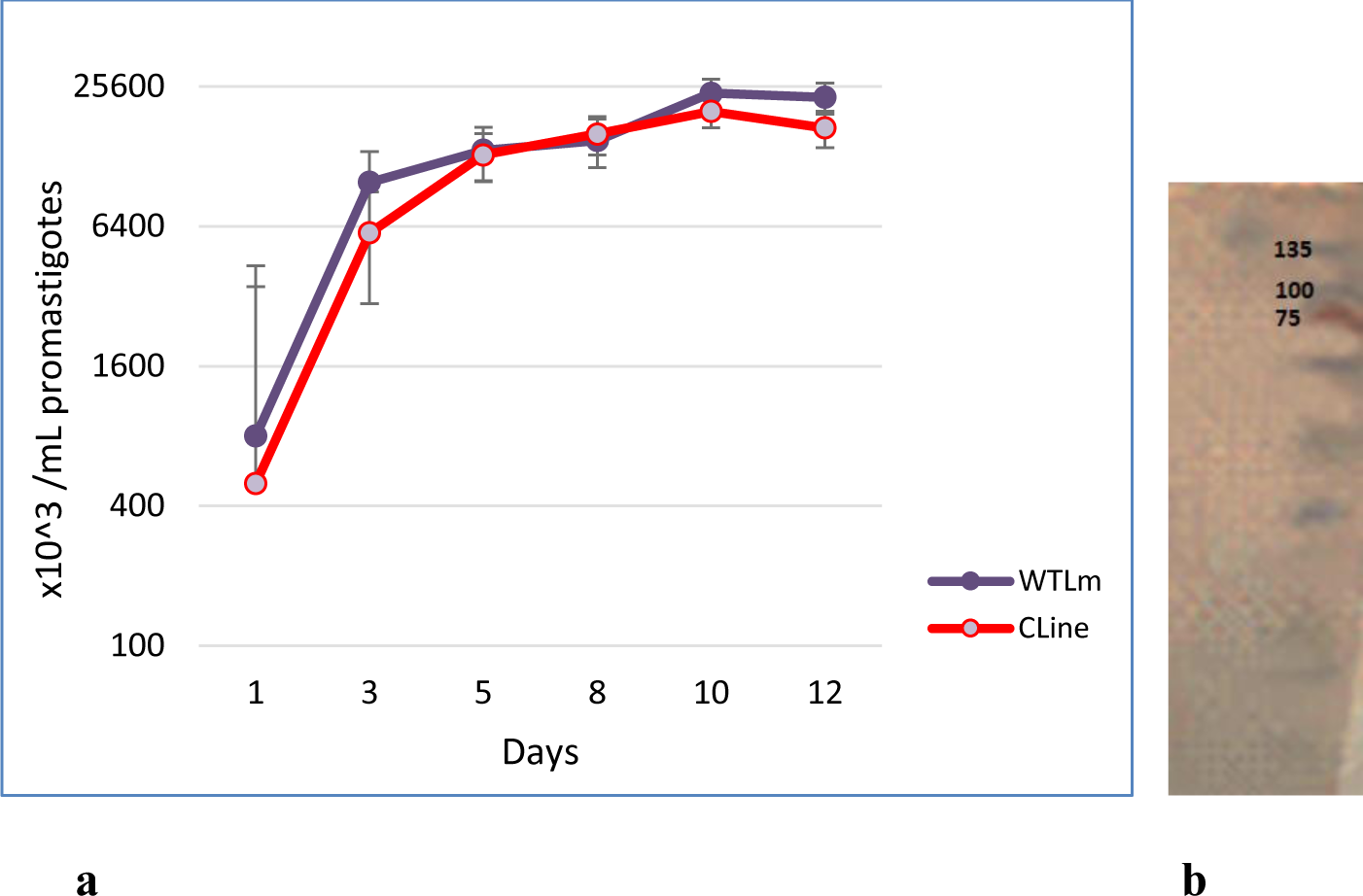
Confirmation of Cas9 and T7 RNAP expression and effect on promastigotes survival. **a)** 5 x 10^5^ promastigotes/ml of WT*Lm* and pTB007 CLine were seeded on a flask separately in M199 medium with supplements and counted daily during 10 consecutive days. Expression of Cas9 and T7 RNAP has no effect on parasite growth as compared to WTLm. **b)** Proteins were extracted from *L. major* CLine expressing Cas9-FLAG (*Lm* pTB007, 162 kDa) promastigote in logarithmic phase. Twenty micrograms was analysed by Western blotting using the anti-FLAG M2 antibody. Protein weight marker in kDa is indicated on the right.

### 3. PCR and Western blot analysis confirmed tagging the c-terminal of UMSBP in *L. major*

To generate transgenic parasites expressing UMSBP-mNG-myc from the endogenous locus, we cotransfected *L. major* pTB007 promastigotes (pTB007 CLine) with a sgRNA cassette targeting the 3′ end of *UMSBP* and a repair cassette containing the puromycin resistance marker and the mNeonGreen-3xmyc tag in frame. The correct integration of the tagging cassette in transgenic promastigotes was confirmed by PCR showing an amplicon of about 920 bp (Figure 3b). The protein was extracted from transgenic strain 2 weeks after culture in the presence of puromycin. Western blot analysis with an anti-myc antibody revealed a band at about 61 kDa corresponding to the expected size of the tagged UMSBP (Figure 3a).

**Fig. 3.**
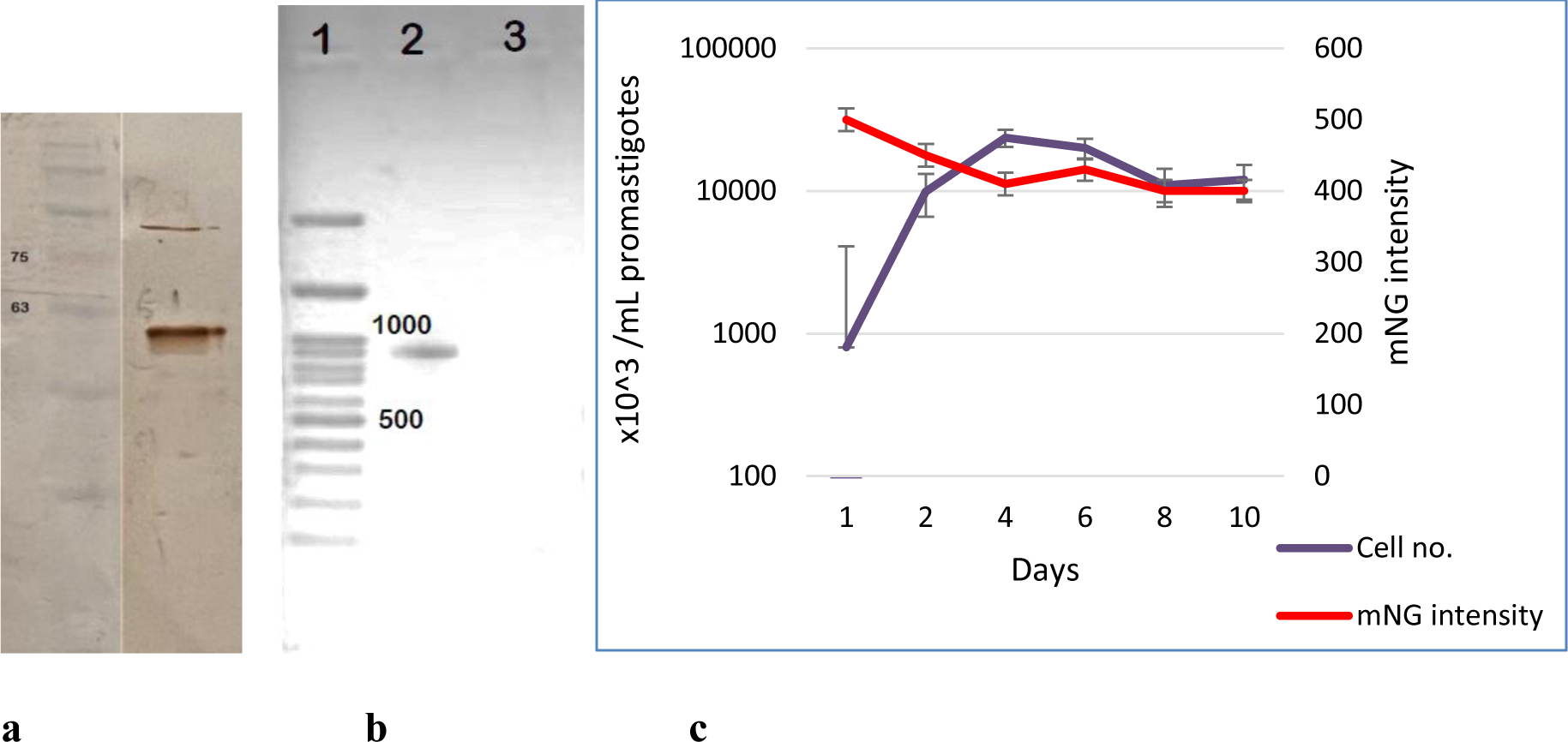
Confirmation of the tagged UMSBP in *L. major*. **a)** The existence of myc fragment in tagged parasite is shown by Western blotting analysis. The protein was extracted from transgenic strain 2 weeks after culture in the presence of puromycin. A 61 kDa band is revealed due to the binding of anti-myc monoclonal antibody to the Myc fragment in the drug resistance cassette, corresponding to the expected size of the tagged UMSBP. **b)** Confirmation of correct integration of the tagging cassette in transgenic promastigotes using specific PCR. The presence of mNG construct in transfected *L. major* parasites were detected. Lane 1 shows 100 KDa ladder, lane 2 shows 920 bp band obtained by using primers as mentioned in table 2. Foreward primer was designed to the coding region of the gene and the reverse primer was designed to the antibiotic resistance marker. Lane 3 is non-template control. **c)** Promastigotes of pTB007 CLine and mNG-tagged strains were seeded separately in M199 medium with supplements on culture flasks and counted daily during 10 consecutive days. Tagging has no effect on parasite growth as compared to pTB007 CLine.

### 4. Constant fluorescence intensity of tagged parasite during promastigote growth

Fluorescence intensity measurements showed that it is constant during promastigote growth and that the expression of the mNG-myc reporter fused to the C-terminus of UMSBP does not lead to any growth defects (Figure 3c). We could not detect the any difference between UMSBP tagged *L. major* and WT*Lm* in analysis by fluorescence microscope probably due to the small size of UMSBP protein integrated in kinetoplast structure.

### 5. Validation of the CRISPR Cas9 gene editing toolkit in generation of mutant *L. major*

Next, we targeted the *UMSBP* locus to generate mutants in a single round of transfection. pTB007 CLine promastigotes were transfected with two sgRNA template targeting upstream and downstream of the UMDBP CDS, and two repair cassettes containing the puromycin and the blasticidin resistance genes. We attempted to delete two allele of UMSBP gene of *L. major* to generate null mutant to determine how this deletion affects the function of the parasites, but we could not obtain parasites that were resistant to both antibiotics and they failed to survive the culture.

WE confirmed the correct integration of the resistance genes at UMSBP locus and generation of *Lm*UMSBP^+/-^ drug-resistant cell population by PCR (Figure 4). We designed 3 primers 1 forward complementary to CDS of UMCBP gene and 2 reverse primers which targeted the antibiotic resistance markers (blasticidin and puromycin). The PCR reactions generated two products, 926 bp for blasticidin and 500 bp for puromycin fragments. Results of sequencing confirmed the correct integration of resistance cassette in the locus.

**Fig. 4.**
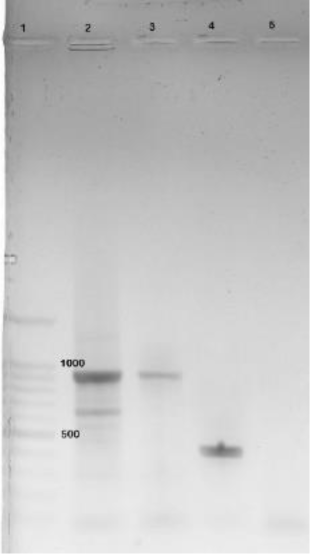
Confirmation of deletion of UMSBP gene. To assess the loss of the target gene in the strains, genomic DNA was isolated from parasites collected after two passages post-transfection. designed 3 primers 1 forward complementary to CDS of UMCBP gene and 2 reverse primers which targeted the antibiotic resistance markers (blasticidin and puromycin) Since we were unable to obtain a parasite that could survive in the presence of both antibiotics, there is no PCR products with two bands corresponding to two antibiotic resistance fragments. Lane 1 shows 100 bp ladder, Lane 2 shows 926 bp of blasticidin resistance cassette, lane 3 shows the same DNA sample upon 1/10^th^ dilution. Lane 4 shows 458 bp of gene locus in WT*Lm* strain.

### 6. The *Lm*UMSBP^+/-^ mutant parasites caused reduced infectivity in cultured macrophages

Due to the role of UMSBP gene in kDNA replication and parasite respiratory chain metabolism, we examined the infectivity, multiplication and growth of *Lm*UMSBP^+/-^ mutant strain compared to WT*Lm* and pTB007 CLine in J774 macrophage cell line (Figure 5).

**Fig. 5.**
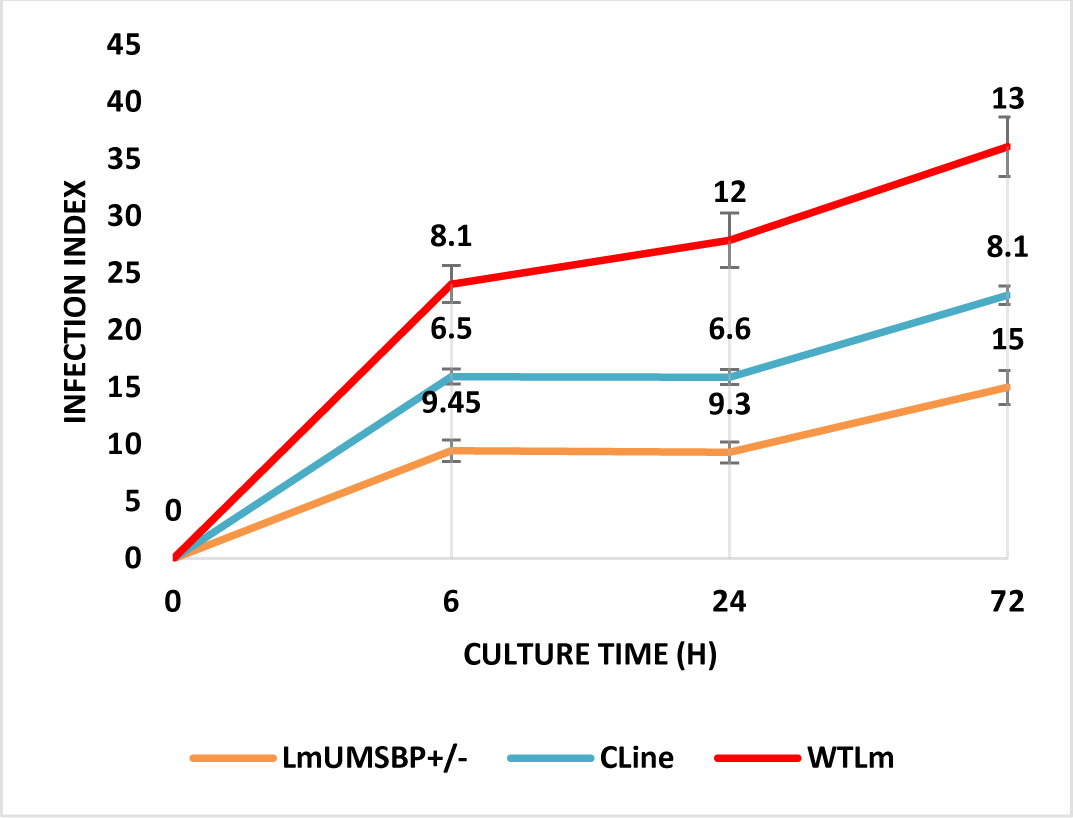
The *Lm*UMSBP^+/-^ mutant cells causes reduced infectivity in cultured macrophages. Stationary phase of *L. major* KO, pTB007 CLine, and WT*Lm* strains were used to infect J774 macrophages. Briefly, 2×10^5^ macrophages were pre-seeded a day in advance in 8-wells chamber slide and after the cells adhered to the slide, *Leishmania* promastigotes were added at a ratio of 10:1 (parasite:macrophage) in 200 μl RPMI 1640 medium and the number of infected cells and internalized parasites were counted in 100 macrophages, at 6, 24 and 72 h after infection. . Results are calculated by multiplying the number of infected macrophages by the number of amastigotes per infected macrophage and expressed as infection index.

Quantitation of infection field in macrophages 6 hours after infection showed that *Lm*UMSBP^+/-^ mutant promastigotes infected a significantly lower number of macrophages (20% of the cells with an average of 3.54 parasites per cell) compared to WT*Lm* (No.=35%, average= 4.17) and pTB007 CLine strain (No.=29%, average= 4.16) (*P*<0.05). Similarly, 24 hours after infection, WT*Lm* and pTB007 CLine strains infected 32% and 24% of the macrophages with 2.43 and 2.4 parasites per cell, respectively, whereas the *Lm*UMSBP^+/-^ mutant strain infected only 16% of the macrophages, with an average number of 1.63 parasites per cell (*P*<0.05). A more pronounced difference was observed after 72 hr of infection, where *Lm*UMSBP^+/-^ mutant strain infected 14% of J774 cell lines with an average number of 1.24 parasites per cell, while WT*Lm* and pTB007 CLine infected 30 and 29% with an average number of 3 and 2.98 parasites per cell, respectively (*P*<0.001).

Therefore, the *Lm*UMSBP^+/-^ mutant promastigotes infected a significantly lower number of cells and proliferated as intracellular amastigotes at a significantly lower rate compared to the control cells (Figure 5).

### 7. Deletion of one allele from the UMSBP gene reduced the growth of the parasite in the culture medium

The growth of the *Lm*UMSBP^+/-^ mutant strain compared to the WT*Lm* strain was evaluated and recorded daily counting during 10 days by microscopic examination. The results showed that parasites that were resistant to both antibiotics failed to survive the culture and null mutant strain (ΔUMSBP) did not show growth trend similar to other strains. Investigating the growth and proliferation of the single KO mutant strain (UMSBP^+/-^) (*i.e.* phenotypes resistant to one antibiotic) showed that the growth of these promastigotes decreased significantly from the early logarithmic phase (*P*<0.05) up to day 10 of growth analysis (Figure 6).

**Fig. 6.**
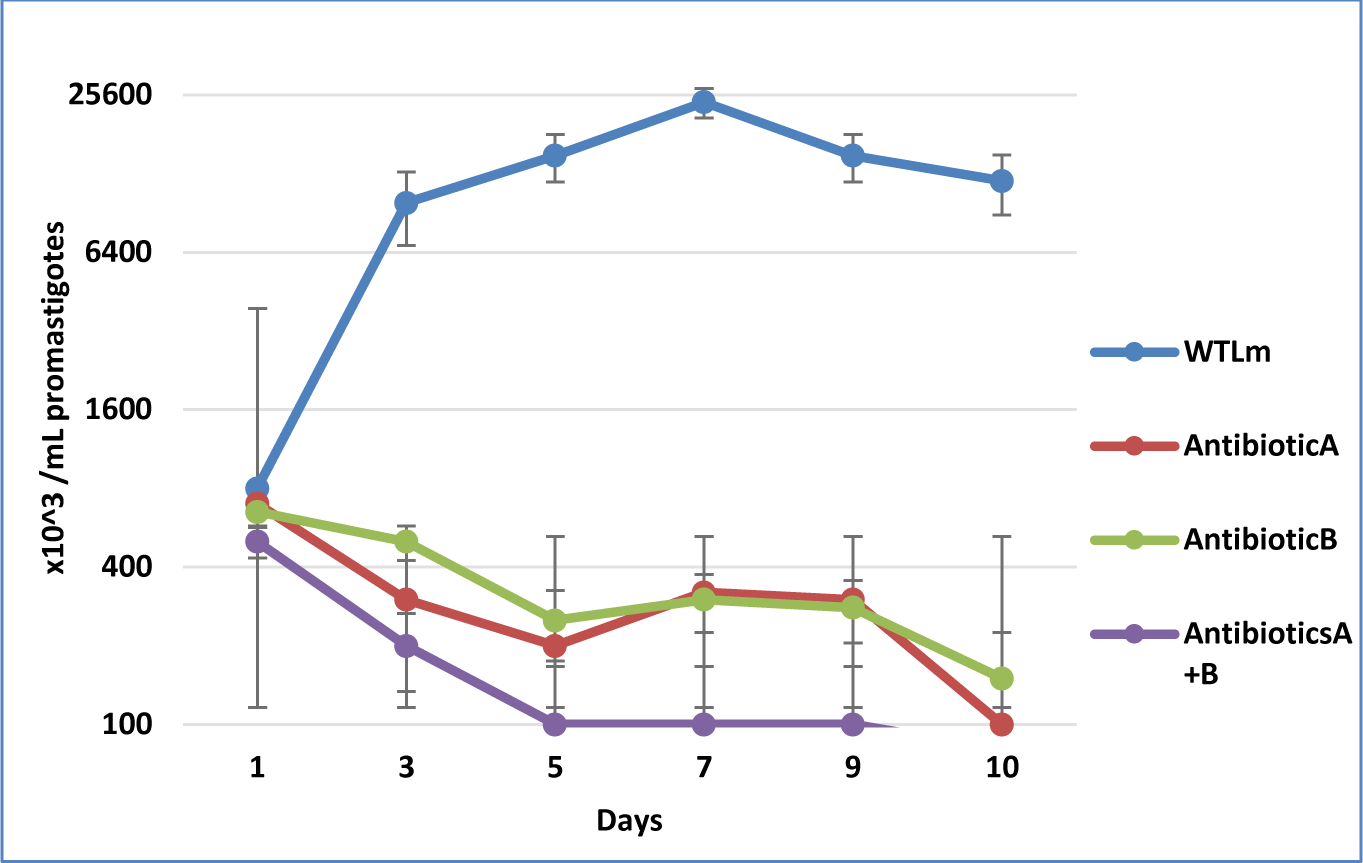
Effect of UMSBP single allele knock out on the promastigotes survival. 5×10^5^ promastigotes/ml of WT and *Lm*UMSBP^+/-^ mutant strains with either of puromycin (A) or blasticidin (B) antibiotic cassettes and ΔUMSBP with both antibiotic cassettes were seeded on a flask separately in M199 medium with supplements and counted daily during 10 consecutive days. Null mutant strain could not survive and the growth of *Lm*UMSBP^+/-^ mutant strain (i.e. phenotypes resistant to one antibiotic) promastigotes decreased significantly from the early logarithmic phase up to day 10 of growth.

### 8. *Lm*UMSBP^+/-^ mutant strain showed increased late apoptosis and necrosis

In healthy cells, annexin-V binds specifically to phosphatidylserine (PS) in the inner layer of the plasma membrane which translocate to the outer membrane after the start of apoptosis. Also, the vital dye propidium iodide (PI) enters only the cells whose plasma membrane structure has been destroyed. Using these characteristics, it is possible to differentiate cells that are in the early stages of apoptosis (Annexin V+/PI-) from those in the final stages. As shown in Figure 7, the three groups WT*Lm*, pTB007 CLine and *Lm*UMSBP^+/-^ have been compared in terms of PI positivity and the amount of PS in the membrane, and it was suggested that deleting one allele of the UMSBP gene in *L. major* parasite significantly increased cells that are in the final stage of apoptosis and necrosis (P<0.05).

**Fig. 7.**
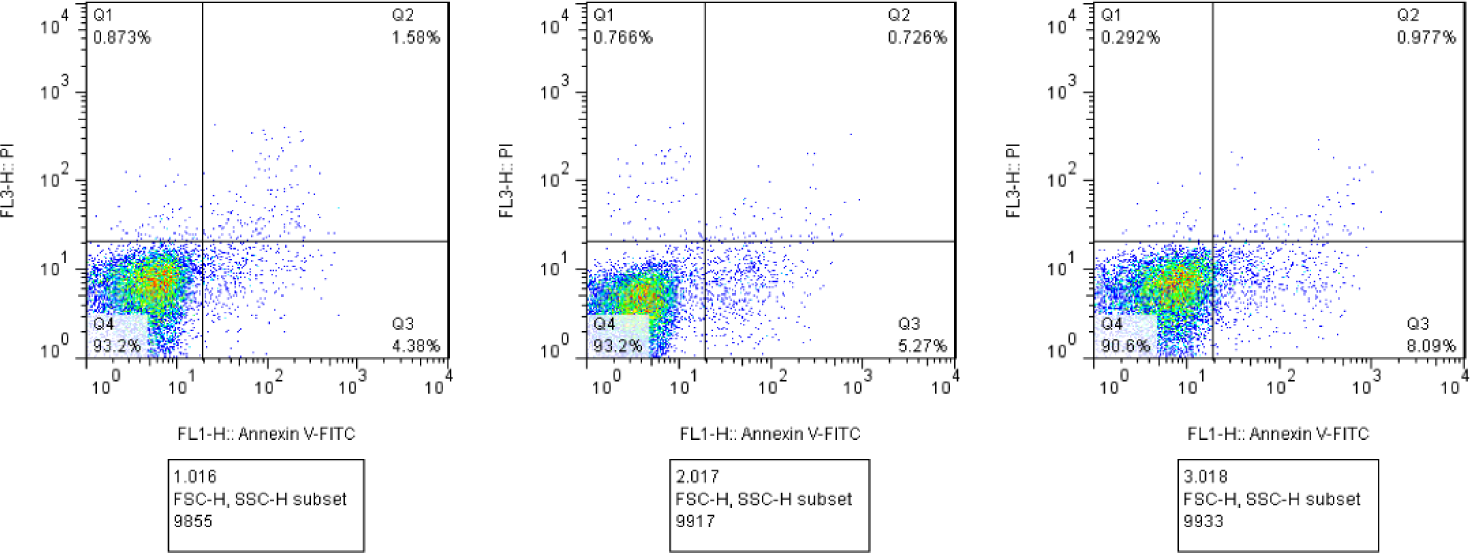
Increased apoptosis and necrosis in *Lm*UMSBP^+/-^ mutant strain. The binding of phosphatidylserine to annexin V in the early stages of apoptosis, late stages and necrosis, in *L. major* WT, CLine and KO strains was measured by the annexin V kit. Fluorescence emission at 530 nm (FL1, Anexin V-FITC) and >575 nm (FL3-PI) were analyzed. According to the flow cytometry reading, deleting one allele of the UMSBP gene in *L. major* parasite significantly increased cells that are in the final stage of apoptosis and necrosis: 1: *Lm*UMSBP^+/-^ mutant 2: Cas9/T7 expressing pTB007 CLine 3: WT*Lm* strains.

### 9. Knockout of the UMSBP gene decreased TXNPx and TryS genes expression during growth of *L. major* promastigotes

In procyclic phase (day 2-3), expression of TXPNx was down regulated in *Lm*UMSBP^+/-^ strain compared to WT*Lm* (about 8.2 folds) (P<0.05) and in stationary phase (day 5-6), expression of TXPNx in *Lm*UMSBP^+/-^ had 16.7 folds decrease compared to WT*Lm* (P<0.001).

On the other hand, expression of TryS, significantly decreased in *Lm*UMSBP^+/-^ strain compared to WT in both procyclic (45.25 folds) (P<0.001) and stationary phase (P<0.001).

### 10. Inoculation of the *Lm*UMSBP^+/-^parasites into BALB/c mice did not cause severe pathology

It was necessary to establish whether single allele deletion of UMSBP gene (*Lm*UMSBP+/−) had any effect on the ability of *L. major* to cause cutaneous infections. To this end, *Lm*UMSBP+/− strain in parallel with WT*Lm* were injected s.c. into the footpad of BALB/c mice and both footpad thickness and parasite burden were measured.

As shown in Figure 8a, swelling of footpad due to WT*Lm* started early, by weeks 3, but *Lm*UMSBP^+/-^ did not induce an early progressive lesion development. *Lm*UMSBP^+/-^ strain failed to produce ulcer in the infected footpad of mice 10 weeks after infection, whereas WT*Lm* did induce observable ulcer by week 6 after infection (Figure 8b). While, the WT*Lm* infected mice developed ulcerative and progressive cutaneous lesions, *Lm*UMSBP+/− infected mice showed a nodular induration and didn’t show any ulcerative lesions at footpad even by the study endpoint (20 weeks).

**Fig. 8.**
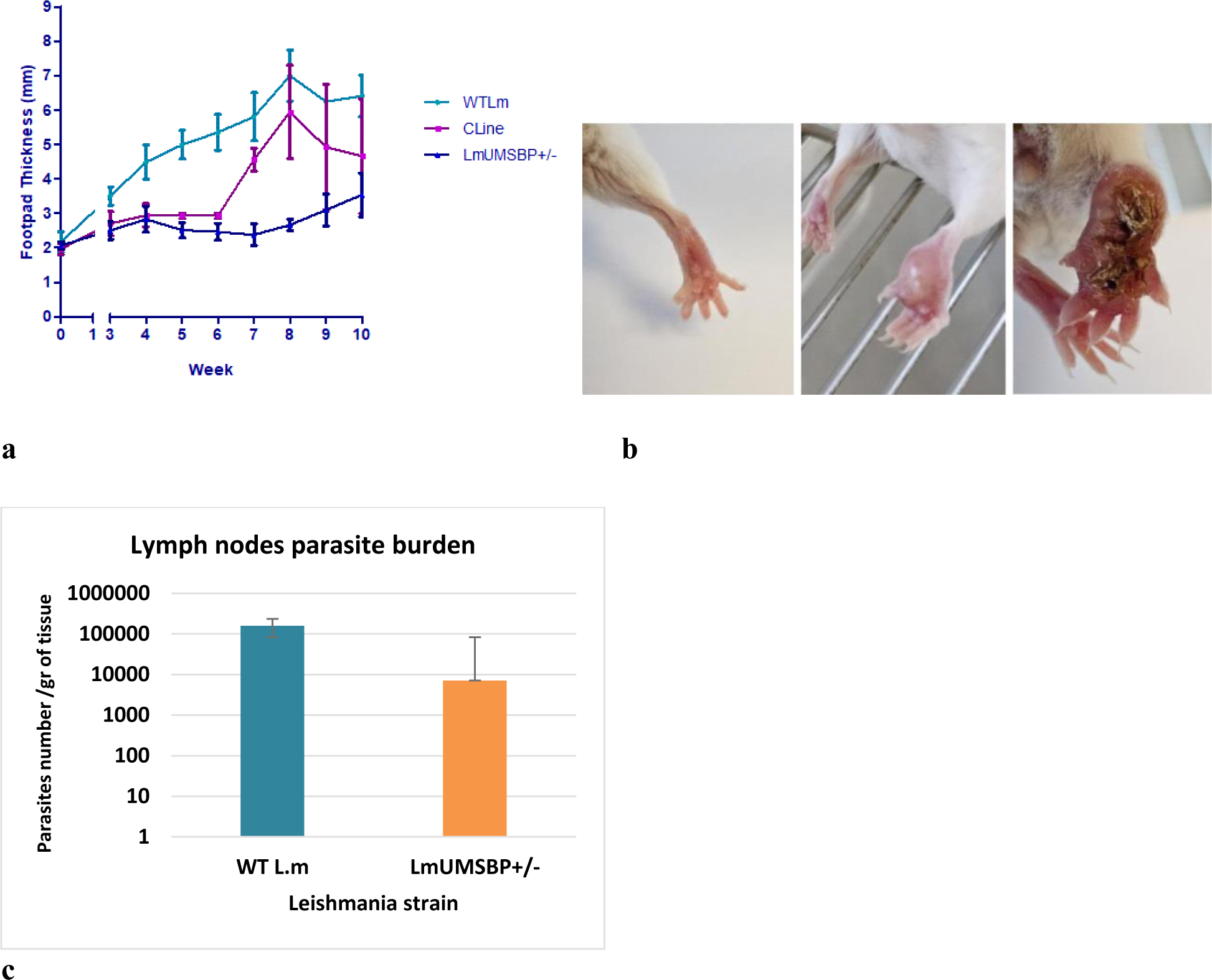
No severe pathology was observed in mice after *Lm*UMSBP^+/-^ injection. 1 × 10^7^ WT*Lm* or *Lm*UMSBP^+/-^ promastigotes in the stationary phase were injected s.c. into the right footpad of mice. **a)** Footpad thickness was measured weekly after infection using a metric caliper up to 10 weeks post infection (wpi). **b)** Representative pictures of mice footpad inoculated with either *Lm*UMSBP^+/-^ at 5 wpi (left) and at 10 wpi (middle) or WT*Lm* (right) promastigotes at 8 wpi. **c)** Parasite burden in the draining lymph nodes at 8 wpi determined by LDA method.

In parasite burden determination, at 8 weeks following infection, the *Lm*UMSBP+/− infected mice had a significantly lower (P<0.05) detectable parasites (approximately 1-2 log fold reduction) as compared to WT*Lm* infected mice that had ∼1.6×10^5^ parasites per g of tissues (Figure 8c). These observations confirm that at 10 weeks post-infection, *Lm*UMSBP+/− is unable to induce severe pathology at the site of injection or visceral organs.

### 11. Immunization with *Lm*UMSBP^+/-^ parasites induced partial protection against live *L. major* challenge in BALB/c mice

To investigate the protective efficacy of *Lm*UMSBP+/− against wildtype *L. major*, susceptible BALB/c mice were immunized with a single i.d. injection with 2×10^6^ stationary phase *Lm*UMSBP+/− in one ear. Four weeks post-immunization (wpi), mice were challenged with 2×10^6^ WT*Lm* in the contralateral ear via the i.d. route (Figure 9). Following challenge with WT*Lm*, lesion development was assessed up to 10 weeks in BALB/c mice. Result showed a significantly higher lesion size as started to increase from week 5-6 post challenge (wpc) (Figure 9a). In the non-immunized (PBS) group, mice developed a non-healing open ulcer that progressively increased in size upon live challenge. No open ulcers were observed in the *Lm*UMSBP+/− immunized group after challenge and only a moderate swelling was seen that subsided from week 9-10 post-challenge (Figure 9b).

**Fig. 9.**
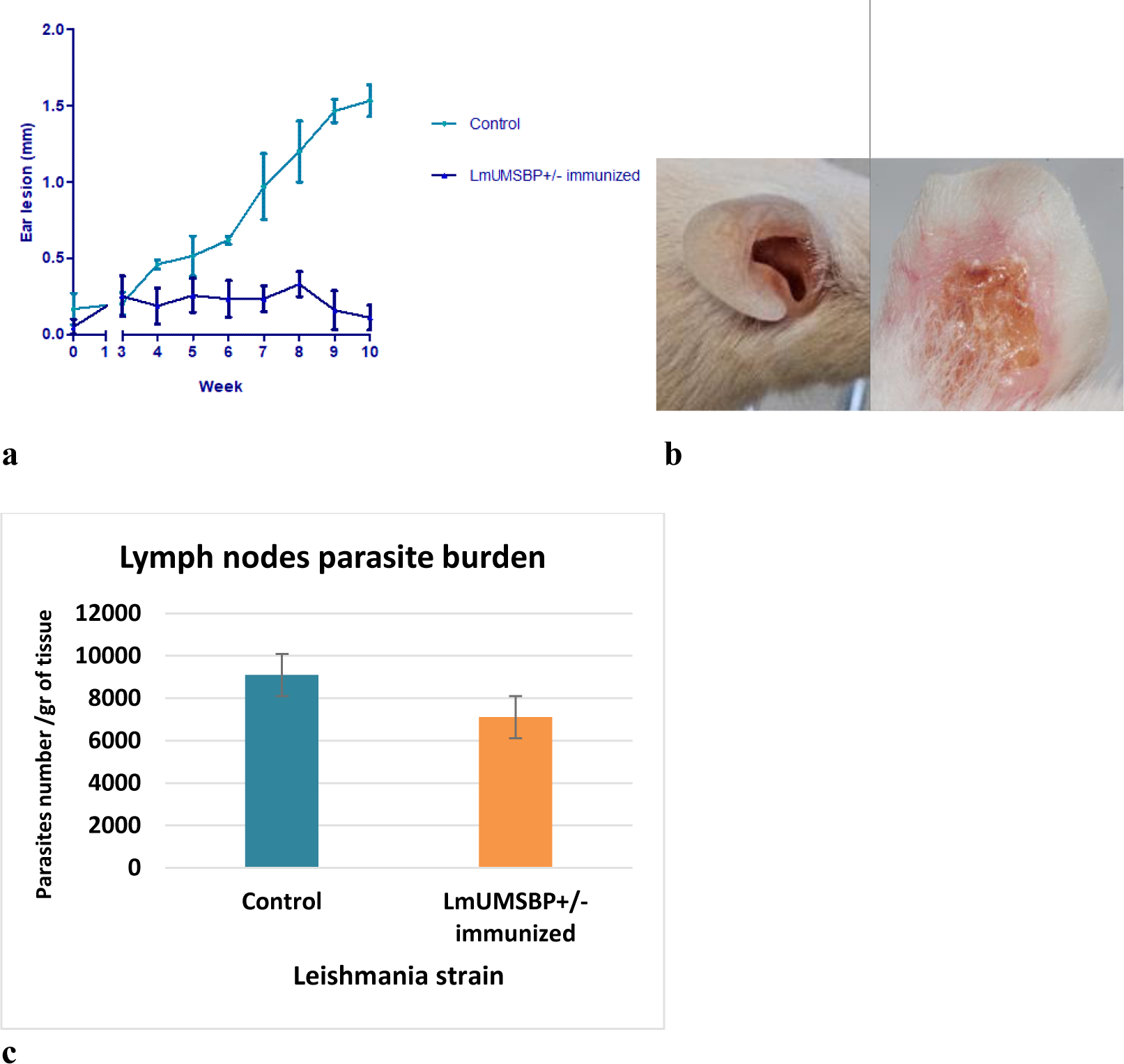
Partial protection against live *L. major* in *Lm*UMSBP+/− immunized mice. For vaccination and immunological studies, two groups of BALB/c mice were immunized i.d. in the left ear with either 2×10^6^ *Lm*UMSBP^+/-^ parasites, or PBS. After 4 weeks post immunization, both groups of mice were challenged i.d. in the contralateral ear with 2×10^6^ live stationary phase WT*Lm* promastigotes. **a)** Ear lesion size was measured weekly after infection using a metric caliper up to 10 weeks post challenge (wpc). **b)** Representative pictures of challenged ear of *Lm*UMSBP^+/-^ immunized (left) or PBS control (right) mice at 10 wpc. **c)** Parasite burden in the draining lymph nodes at 10 wpc determined by LDA method.

The parasite load of draining lymph nodes were also quantified at 10 wpc revealing that the immunized group had a significantly lower parasite load compared to the non-immunized control group (P<0.05) (Figure 9c).

### 12. Immunization with *Lm*UMSBP^+/-^ parasites induced a Th1-dominant immune response against *L. major* challenge in BALB/c mice

Having shown above that *Lm*UMSBP+/− induces partial protection against needle challenge, we compared the immune response between *Lm*UMSBP+/− immunized group (4 wpi) and PBS control group. We titrated IFN-γ and IL-12 as Th1 type cytokines, and Il-4 and Il-10 as Th2 type cytokines in the supernatant of lymph nodes cell culture upon *L. major* antigen (SLA) stimulation (Figure 10). Data showed a significantly higher IFN-γ (P<0.001) and significantly lower IL-4 (P<0.001) levels in cells from *Lm*UMSBP+/− immunized mice compared to cells from PBS control group in response to SLA. The levels of IL-12 and IL-10 showed no significant changes in cells from *Lm*UMSBP+/− immunized mice compared to cells from PBS control group (Figure 10).

**Fig. 10.**
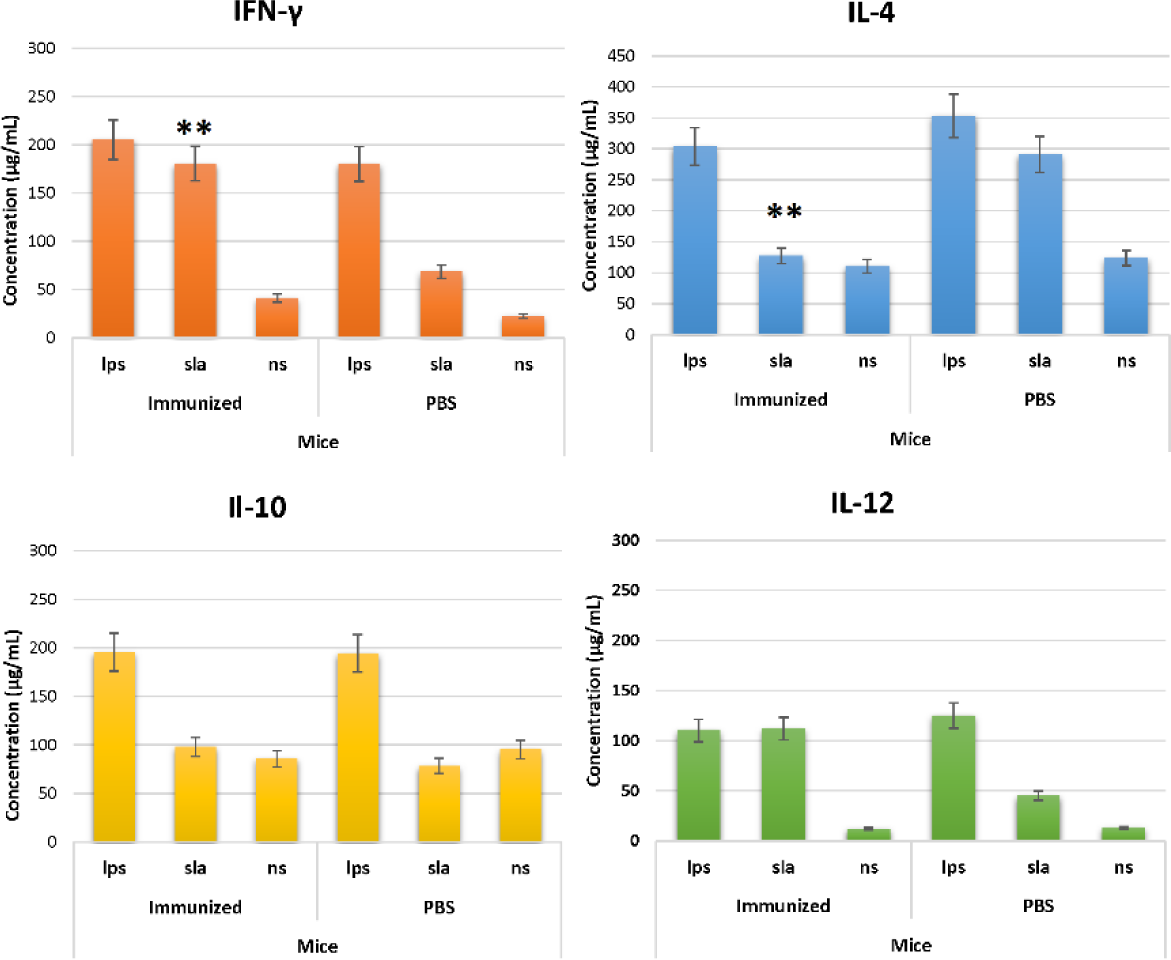
Titration of cytokines in cell culture supernatant of immunized mice. Four weeks after mice immunization, immunized and PBS control mice were sacrificed and lymph nodes were isolated and resuspended in complete RPMI 1640 medium at a density of 2×10^6^/mL in 96-well plates in the presence of either LPS, SLA or without stimulation (ns). Plates were incubated at 37 °C with 5% CO2 in humidified atmosphere, and the levels of IL-12, IFN-γ, IL-4 and IL-10 were titrated on 72-h culture supernatants using sandwich ELISA method. Data shows mean ±SD of each group of mice (*n*=5) for each stimulation.

## Discussion

CRISPR/Cas9 is an RNA guided efficient genome editing tool to delete or disrupt genes, generate point mutations, and add tags to endogenous genes using endonuclease derived from the bacterium *Streptococcus pyogenes* (19). Due to its simplicity, versatility, and high efficiency, it has been widely used for genome editing in a variety of organisms including the protozoan parasite *Leishmania* (10). To assess the efficiency of a CRISPR/Cas9 gene editing toolkit in *L. major*, we performed C-terminal tagging and the generation of KO mutants on a kDNA-associated gene, UMSBP, which initiates kDNA replication by binding with a conserved universal minicircle sequence (UMS) of kDNA. Previous experiments in *L. donovani* showed *Ld*UMSBP regulates leishmanial mitochondrial respiration and pathogenesis and its deletion leads to kDNA loss and apoptosis like death of promastigotes (20). Interestingly, single knockout of UMSBP in *L. donovani* has no effect on promastigotes survival. However, in intracellular amastigotes demonstrate loss of mRNA level of cytochrome-b and reduced production of ATP. This process interfere with the oxidative-phosphorylation and thereby completely inhibit the intracellular proliferation of *L. donovani* amastigotes in human macrophages and in BALB/c mice (20). We sought to replicate these phenotypes in *L. major* using the CRISPR/Cas9 gene editing strategy to validate the toolkit in this parasite species and to generate a live attenuated parasite which show impairment in amastigote replication stages without the defect in promastigote fitness in culture. The key advantages of the used CRISPR/Cas9 system are that it does not require any gene-specific cloning procedures or in vitro transcription prior to transfections, making it rapid, scalable, and economical (21). We first generated a *L. major* cell line which expresses Cas9 nuclease and T7 RNA polymerase (RNAP) constitutively. The expression of Cas9 was confirmed by Western blot analysis which showed this expression did not alter the growth of *L. major* promastigotes, similar to what has been shown previously with *L. donovani* and *L. mexicana* promastigotes (22).

Present data indicate that the tagging of UMSBP using CRISPR Cas9 was successful in *L. major*, as shown by PCR and fluorescent analysis. This approach is simple and fast and will greatly improve the way we study *Leishmania* genes compared to previous methods of gene tagging (e.g., by episomal expression of fluorescent tag), since part of their endogenous regulation may be better maintained through the conservation of either the 5′ or 3′ UTR. Furthermore, results indicate that UMSBP has been successfully deleted in the whole population, without the need for subcloning as previously reported with other genes using the same CRISPR approach (22). The generation of homozygous ΔUMSBP *L. major* parasites was not successful, indicating that UMSBP gene is probably essential for promastigote survival.

These data validate the use of the CRISPR Cas9 toolkit developed by Beneke et al. in *L. major*. In addition to flexibility, this approach has some other advantages over other developed CRISPR methods. The ability to generate homozygote KO in one single transfection will minimize the complexity and possible phenotypic changes of the transgenic parasites. Furthermore, previously we have found that *L. major* is particularly sensitive to successive *in vitro* culture and may lose its virulence after more than 10 consecutive passages in culture to become unable to infect mice (23). Thus, minimising time in culture is a prerequisite for virulence studies and the current CRISPR/Cas9 method will enhance our ability to manipulate *L. major* parasites.

Mitochondrial redox regulating enzymes, tryparedoxin and TXNPx regulates the oxidation and reduction status of UMSBP and thereby its UMS binding activity (24). In a previous study in *L. donovani,* it was suggested that increase in TXNPx is associated with decrease in the relative UMS binding capacity, when the concentration of tryparedoxin is constant (20).

We observed a noticeable growth defect in *Lm*UMSBP^+/-^ promastigotes, especially at later stages of parasite growth (stationary phase), where dividing procyclic promastigotes undergo a terminal differentiation into infective metacyclic stages. The defect in promastigote growth within culture is a challenge as it indicates that deletion of the gene could potentially affect the parasite’s fitness in culture, posing a complication to the large-scale production of *Leishmania* parasites.

Live parasites have the advantage of the better efficacy to induce long-term protective immune responses as compared to killed parasite vaccines; however, the obstacle to using a live vaccine is the potential risk of disease development which might be overcome by engineering a second generation live-attenuated parasite which can confer protection without associated pathology (25) (26). Therefore, it is believed that generation of attenuated parasites through targeted disruption of genes coding for virulence or optimal growth factors is effective approaches for vaccine development against leishmaniasis (27). However, several challenges must be addressed before the effectiveness of live attenuated vaccines can be evaluated in clinical trials, including concerns related to safety, genetic stability, delivery system and type of memory development (28). In this study, a live attenuated *Lm*UMSBP^+/−^ parasites was produced which elicited protective immunity in susceptible BALB/c mice against *L. major* infection. However, production of null mutant was not successful probably because the gene function is necessary, and we have then evaluated the characteristics of single allele deleted *Lm*UMSBP gene mutant strain throughout the experiment. This could be a challenge due to the potential risk of reverting the single knockout parasites to pathogenic wild type over long time *in vivo*.

In this study, we have compared immune response induced by immunization with *Lm*UMSBP^+/−^ parasite compared with PBS against live *L. major* challenge. Our results demonstrate significant induction of a Th1 response with high IFN-γ and low IL-4 cytokines production in infected BALB/c mice. It is shown that IL-12-driven, IFN-γ dominated Th1 response promotes healing and parasite clearance in *Leishmania* infection in animal models (29). In murine cutaneous leishmaniasis elicited by *L. major* promastigotes the resolution of the skin lesions requires IFN-γ-producing CD4^+^ T helper cells, which develop in the presence of IL-12 and the signal transducer and activator of transcription (STATs) (30, 31). Evidence shows both antigen-specific IFN-γ-producing CD4+ and CD8+ T cells are capable of promoting partial protection against progressive *Leishmania* diseases (13). IFN-γ and iNOS, which converts arginine into citrulline and NO, turned out to be the key effector pathway of macrophages against the intracellular *Leishmania* stage (amastigotes) during the acute phase of infection in different models of *L. major* infection (32, 33). Leishmanial antigens that predominantly stimulate Th1 responses in spleen or lymph node cells from mice infected with *L. major* have commonly been accepted as "potential protective antigens" and therefore promising vaccine candidates. Although, evidence exist which challenge the concept of Th1/Th2 paradigm to use Th1 cells from infected hosts as readouts for antigen selection in vaccine development, however, it is generally well accepted that a Th1 response is essential for protection against experimental leishmaniasis (34, 35).

The *Lm*UMSBP^+/-^ mutant promastigotes showed significant reduction in infection of macrophages and intracellular replication as intracellular amastigotes compared to the WT parasites. While we need parasite growth within the host cells to elicit the effector immune responses and development of memory T cells, this growth needs to be limited to guaranty parasite persistence without pathology (36). Two months following infection, the *Lm*UMSBP^+/−^ parasites were not completely cleared from the spleen and lymphnodes, even a significantly reduction of the parasite load was observed compared to WT*Lm* infected mice. The residual parasite burden observed in mice organs in the *Lm*UMSBP^+/−^ may be important for maintaining long term protection as was reported in other candidate vaccine formulations in animal models (37, 38). However, unlike leishmanization which involves inoculation of virulent parasites that causes lesions at the site of injection, immunization with *Lm*UMSBP^+/−^ parasites seems to be safe as demonstrated by the absence of visible ulcerative lesions in susceptible mice, in spite of persistence of a low number of parasites at the site of inoculation.

In this experiment, prior immunization with *Lm*UMSBP^+/−^ confers partial protection against *L. major* needle challenge in mice as shown by significant decrease in parasite burden and lesion development up to 10 wpc. While normally 2×10^6^ *L. major* parasites cause development of a cutaneous lesion that progressively increases in size, leading to disseminated infection in BALB/c mice, in clear contrast, no open ulcers were developed in the *Lm*UMSBP^+/−^ immunized group and only a moderate swelling was observed. These observations of protective effect of *Lm*UMSBP^+/−^ strain using a cutaneous model of *Leishmania* infection are consistent with previous findings in *L. donovani* species that single allele UMSBP gene deleted parasites could confer protection against visceral leishmaniasis in animal model as a live attenuated vaccine candidate (20).

## Conclusion

In this study, we validated the CRISPR/Cas9 toolkit for *L. major* targeting a kinetoplast-related gene to generate knockout and tagged parasites. The fact that only one single transfection without cloning is required to obtain knockout mutants, makes this approach particularly feasible and suitable for virulence studies, as it prevents possible loss of virulence that may occur during successive *in vitro* culture.

Leishmanization with wild type *L. major* has so far been the only successful immunization strategy for human leishmaniasis, but it has been discontinued due to safety concerns associated with administration of a live virulent organism. Vaccination with attenuated UMSBP^+/−^ *L. major* strain did not cause progressive lesions but retains the ability to provide immunological protection against experimental needle *L. major* infection, so future extensive studies are required to establish whether vaccination with this strain is definitely safe and induce protection in mice and humans leishmaniasis.

## Acknowledgments

We would like to express our sincere gratitude to Olivier Leclercq, Najma Rachidi and Gerald Spaeth from Unité de Parasitologie Moléculaire et Signalisation Département de Parasitologie et Mycologie, Institut Pasteur de Paris for organizing the training program on CRISPR/Cas9 technique.

